# Quinazoline-quinoline bisubstrate inhibitors target eukaryotic translation initiation factor 3 in *Plasmodium falciparum*

**DOI:** 10.1101/2022.12.10.519887

**Authors:** Irina Dobrescu, Elie Hammam, Jerzy M. Dziekan, Aurélie Claës, Ludovic Halby, Peter Preiser, Zbynek Bozdech, Paola B. Arimondo, Artur Scherf, Flore Nardella

## Abstract

Malaria drug resistance is hampering the fight against the deadliest parasitic disease affecting over 200 million people worldwide. We recently developed quinoline-quinazoline-based inhibitors (as compound **70**) as promising new antimalarials. Here we aimed to investigate their mechanism of action by using Thermal Proteome Profiling (TPP). The eukaryotic translation initiation factor 3 (EIF3i) subunit I was identified as the main target of the inhibitor in *P. falciparum*. This protein is not a known drug target in malaria parasites. *P. falciparum* parasite lines were generated expressing either a HA tag or an inducible knockdown of the PfEIF3i gene to further characterize the target protein. PfEIF3i was stabilized in presence of the compound **70** in a cellular thermal shift-western blot assay, confirming that PfEIF3i is a target of quinoline-quinazoline-based inhibitors. In addition, PfEIF3i-inducible knock-down blocks intra-erythrocytic development in the trophozoite stage indicating that it has a vital function. We show that PfEIF3i is mostly expressed in late intraerythrocytic stages and localizes in the cytoplasm. Previous mass spectrometry reports show that EIF3i is expressed in all parasite life cycle stages. Hence, quinoline-quinazoline-based inhibitors allowed to identify PfEIF3i as a valuable target for the design of new antimalarial drugs active all along the life cycle of the parasite.

## Introduction

*P. falciparum* is the malaria parasite responsible for the majority of the 619,000 deaths and the 247 million cases worldwide reported in 2021^1^. *P. falciparum* infection can cause severe clinical symptoms such as anemia, respiratory distress and, in the most complicated cases, cerebral malaria. Parasite multi-drug resistance to the actual treatments, consisting in artemisinin-based combination therapies (ACTs), has emerged in South-East Asia causing first line cure failures^2^. Our group recently developed quinoline-quinazoline-based inhibitors that are fast-acting antimalarials able to kill ring stage parasites and resistant field isolates with a promising *in vivo* activity in a mouse malaria model^3^. We have shown that these compounds reduce the activity of DNA methyltransferases (DNMT) to methylate DNA cytosine in human cells as well as in *P. falciparum* protein extracts. However, the only annotated DNMT in *Plasmodium*, DNMT2, is dispensable in asexual blood stages^4^ making DNA cytosine methylation not likely being the primary drug target of these compounds. Thus, the target protein of the compounds remains to be identified.

In this study, we explored the quinoline-quinazoline inhibitors mode of action in malaria parasites by using the Thermal Proteome Profiling (TPP)^5–7^, in which the proteome in the presence and in the absence of the compound is analyzed by mass spectrometry at different temperatures. This allows to identify the proteins that are stabilized by the compound at increasing temperatures. The assay is based on the principle that, the stability of a protein changes upon interactions with a ligand, such as drugs^8^. TPP employs pulse thermal challenge to denature unstable protein subsets and quantifies remaining soluble protein abundance to identify proteins thermally-stabilized by the drug. Using this technique, we found the eukaryotic translation initiation factor 3 subunit I (EIF3i) as the main protein stabilized by the lead bisubstrate inhibitor **70**^3^.

Eukaryotic translation initiation factors mediate the assembly of tRNA, 40S, and 60S ribosomal subunits into an 80S ribosome at the initiation codon of mRNA, a complex process required for protein synthesis initiation^9^. Among these factors, the largest complex EIF3, comprising 13 subunits^10^, has a crucial role in different steps of translation initiation and is involved in termination and ribosomal recycling. Due to its multicomplex network, EIF3 is a major actor in many biological processes and can be associated to functional disorders^11,12^. Its dysregulation is linked to different types of cancer (reviewed in ^13^), and elevated levels of the EIF3i subunit are found in human cancers, suggesting an important role cell proliferation regulation^14^ and cell cycle differentiation. EIF3i has not been studied in *Plasmodium*, we therefore explored the essential functions of this protein in *P. falciparum* blood stage development to better understand its potential as a novel antimalarial drug target.

## Results

### Identification of compound 70 putative targets by TPP

To identify the target(s) of the bisubstrate inhibitor **70**^3^ in *P. falciparum*, parasite protein extracts were treated with increasing concentrations of the compound and fractions were exposed, in parallel, to different temperatures at 37°C (non-denaturing control), 51°C and 57°C. The soluble protein abundance in presence of varying drug concentrations was quantified by mass spectrometry, relative to the vehicle control sample. Four proteins were found to exhibit significant (≥3*Median Absolute Deviation in the Δ Area Under the Curve) drug-concentration dependent stabilization (Fig. 1A), including the eukaryotic translation initiation factor 3 subunit I (EIF3i), 40S Ribosomal Protein S15, Zinc Finger Protein and 3-hydroxyisobutyryl-CoA hydrolase. Among those, EIF3i exhibited the strongest stabilization achieving nearly 3-fold stabilization at the highest drug concentration under the 51°C thermal challenge condition, with effective drug concentration of 1.5µM to evoke stabilizing response (Fig. 1B). The presumed target of compound **70**, the PfDNMT2 was not detected in the assay.

**Figure 1.**
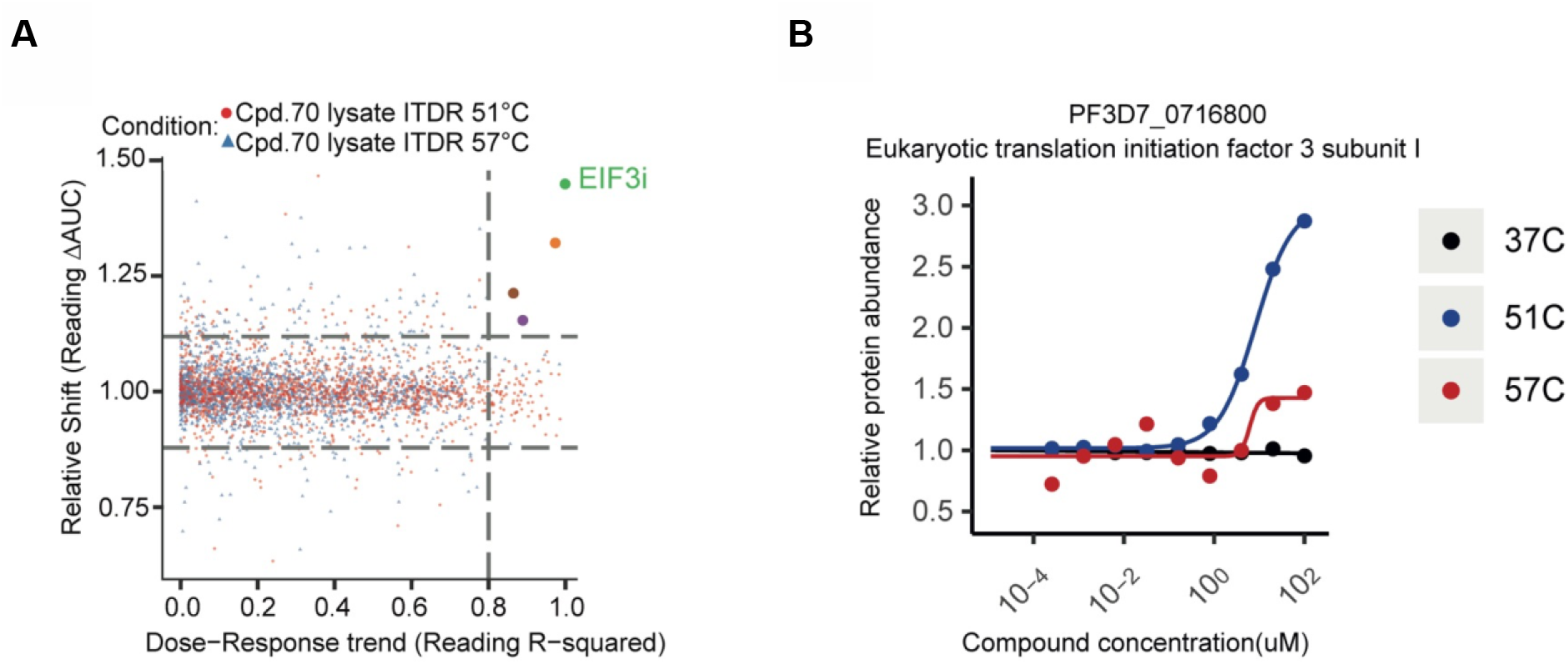
PfEIF3i is stabilized by compound 70. Thermal Proteome Profiling Protein target engagement by compound **70. (A)** Whole proteome analysis in lysate ITDR (isothermal dose response) experiments under 0-100 µM of compound **70** treatment with thermal challenges at 51°C (blue triangles) or 57°C (red circles). Distribution of protein stabilization is plotted as a function of R^2^ value (goodness of curve fit) against ΔAUC (area under the curve of heat-challenged sample normalized against nondenaturing 37°C control) for all proteins detected in the assay. Three times of median absolute deviation (MAD) of ΔAUC in each dataset (MAD × 3) and R^2^ = 0.8 cutoffs are indicated on the graph. Significant hits are highlighted in other colors. **(B)** Thermal stabilization profile of EIF3i identified in **(A)**. Stabilization under thermal challenges [51°C (blue) or 57°C (red)] is plotted relative to no-drug control with non-denaturing control (37°C) in black.

### Compound 70 stabilizes PfEIF3i

To validate the results obtained in the thermal proteome profiling (TPP) analysis, a transgenic *P. falciparum* NF54 parasite line was generated by introducing a 3x hemagglutinin epitope tag (3xHA) expressed in frame with *eif3i* (Fig. 2A). After verification of the correct genomic integration and HA-tagging of EIF3i (Fig. 2B), we used this transgenic parasite strain in the Cellular Thermal Shift Assay (CETSA)^6^ analysis followed by western blot (CETSA-WB). In this assay, parasite protein extracts were treated with 30µM of compound **70** and exposed to different temperatures and analyzed by western blot to assess the level of the tagged EIF3i protein. Upon increase of the temperature, the EIF3i protein is stabilized when treated with the compound **70** compared to the DMSO control (Fig. 2C). The protein stabilization occurs from 40°C until 52°C (Fig. 2D), in agreement with the TPP assay in which the best EIF3i stabilization is observed at 51°C (Fig. 1B). At higher temperatures we observe no difference in protein quantity between DMSO and compound **70** treatment. In addition, we observe no protein stabilization when PfEIF3i-HA parasites are treated with pyrimethamine (negative control) at any of the used temperatures (Fig. 2E, 2F).

**Figure 2.**
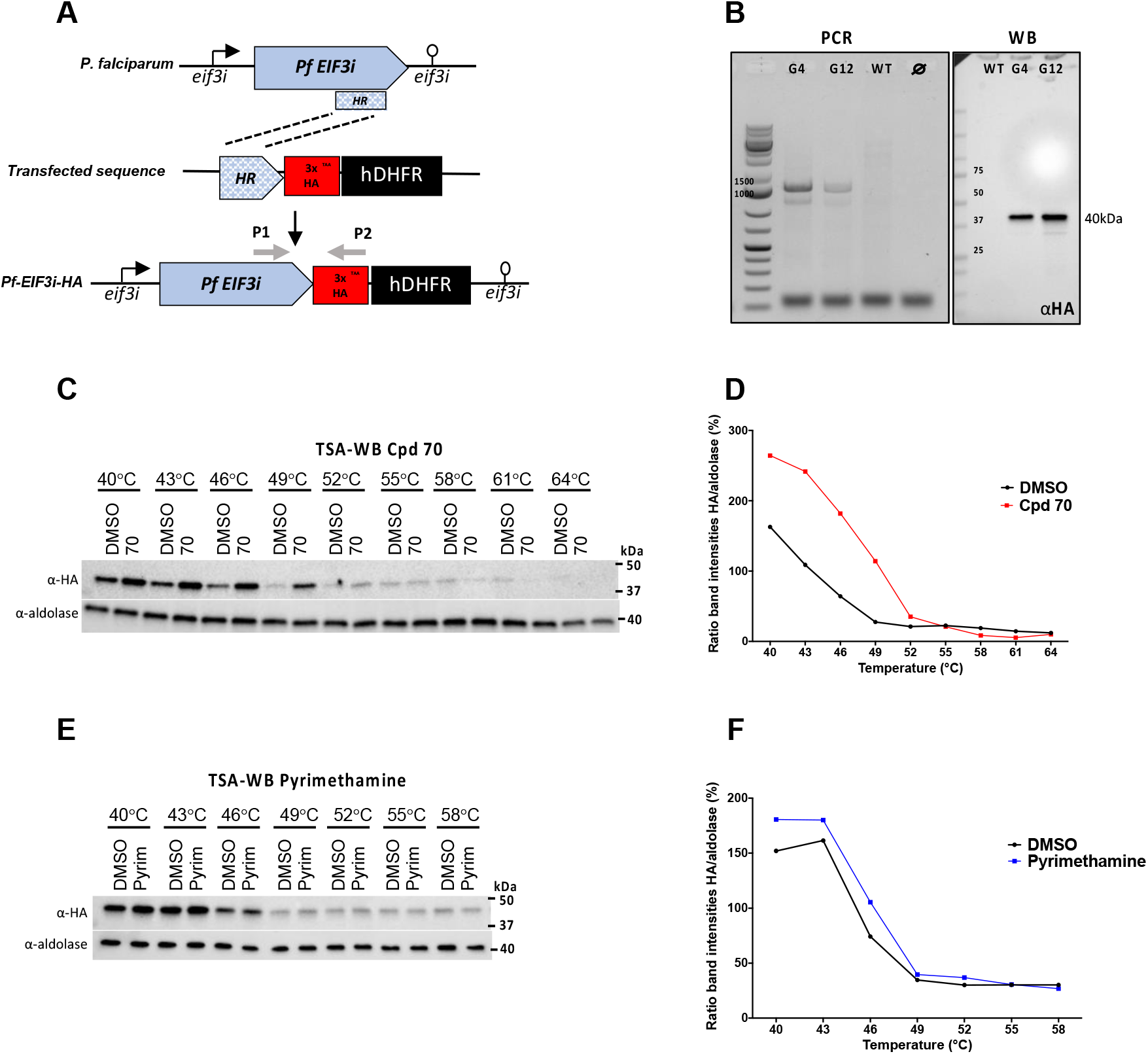
Western blot detection of PfEIF3i stabilization in parasite extracts treated with compound 70. **(A)** Strategy to generate PfEIF3i-HA parasite line by tagging EIF3i with 3xHA tag at the end of the coding sequence (cds). HR, homology region, arrow boxes, coding sequences; arrows, 5′UTR; open circle lollipop, black box hDHFR selection marker cassette; red box, 3xHA-tag; green box; grey arrows P1 and P2, primers used for the PCR. **(B)** After transfection, the integration was validated by PCR and HA expression was confirmed by western blot using anti-HA antibodies. **(C, D)** PfEIF3i-HA extracts were collected, treated with 30 µM of compound **70** for 1h at RT and exposed to a temperature gradient (40-67°C). Soluble protein levels were run in a SDS-PAGE gel and EIF3i levels were quantified by western blot using anti-HA antibodies; anti-aldolase antibodies were used as loading controls. Band intensities were measured with image lab software (Biorad) and graphs represents the HA signal normalized to the aldolase band intensities in percentage. Results were obtained in two independent experiments; figures are representative of one experiment. **(E, F)** Pyrimethamine was used as a negative control in the same conditions as for compound **70**.

### Inducible PfEIF3i knock-down

As we confirmed that PfEIF3i is a target of compound **70**, a fast-acting antimalarial agent active in resistant strains, we studied the function of PfEIF3i. To this aim, we generated a second transgenic parasite PfEIF3i-HARibo line that possess an HA-tag in frame with *eif3i* and an additional *glmS* ribozyme sequence (Fig. 3A) allowing the conditional knock-down of the protein upon glucosamine addition^15^. After transfection, selection and cloning, integration of the sequences was confirmed by PCR using specific primers (Fig. 3B). The two selected positive clones show correct HA expression detected by western blot at the size of EIF3i (37 kDa) (Fig. 3B).

**Figure 3.**
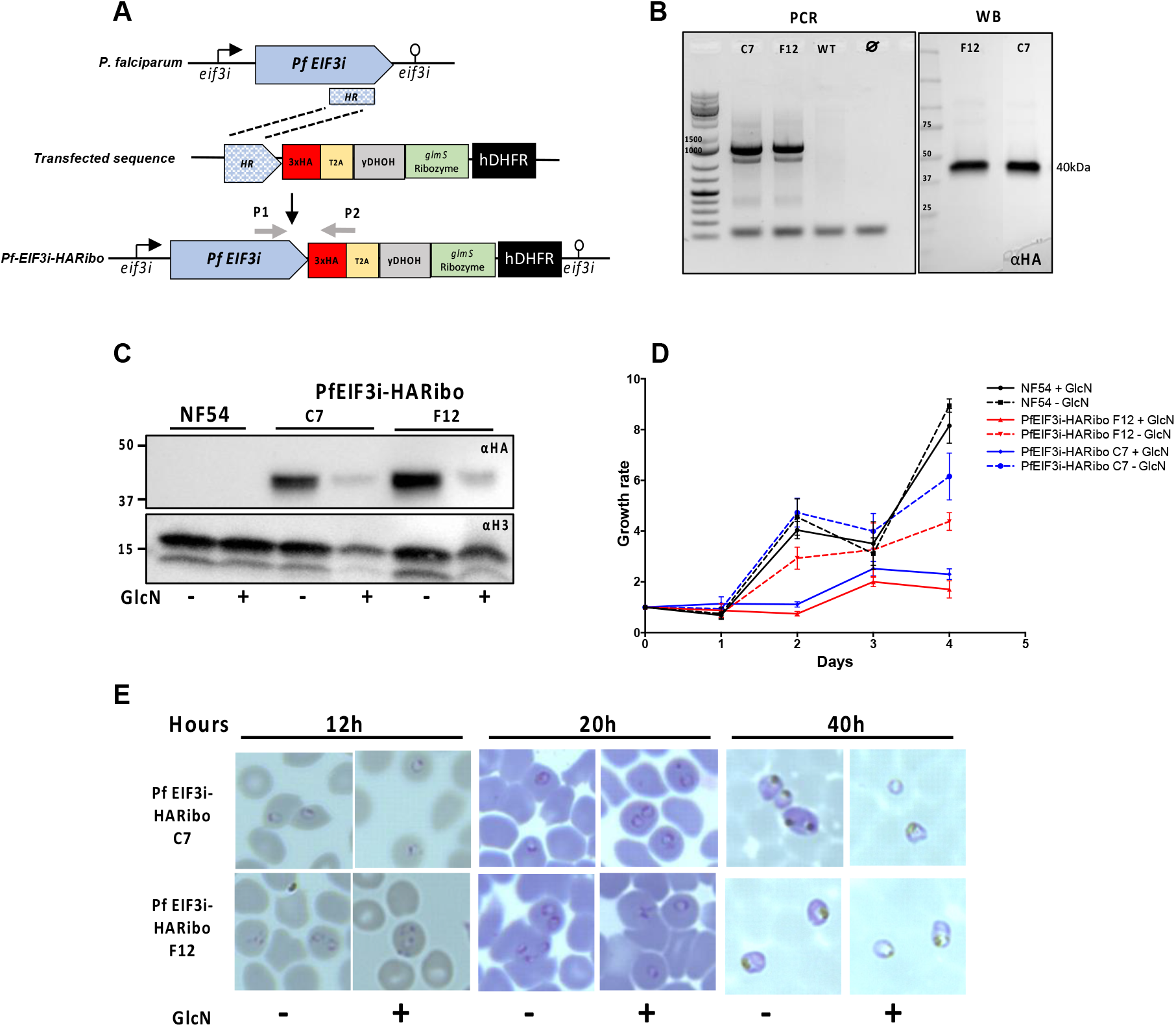
Addition of Glucosamine efficiently knocks-down PfEIF3i protein levels and impact growth. **(A)** Strategy to generate PfEIF3i-HARibo parasite line with 3xHA tag followed by a *glmS* ribozyme sequence. HR, homology region; arrow boxes, coding sequences; arrows, 5′UTR; open circle lollipop, 3’UTR; black box hDHFR, selection marker cassette; red box, 3xHA-tag; green box, *glmS* ribozyme cassette; yellow box, T2A peptide; grey box, yDHOH cassette; grey arrows P1 and P2, primers used for the PCR. **(B)** After transfection, the integration was validated by PCR and correct HA expression was confirmed by western blot using anti-HA antibodies. **(C)** Extracts of PfEIF3i-HARibo clones C7 and F12 were collected after saponin lysis from parasite cultured with or without glucosamine (GlcN) for one cycle. EIF3i protein levels were quantified by western blot using anti-HA monoclonal antibodies and anti-H3 antibodies as loading controls. **(D)** Parasite were cultivated in presence or not of GlcN (added at day 0, D0) and growth was assessed for 4 days, every 24h, by flow cytometry using SYBR Green I staining. **(E)** Synchronized parasites were cultivated in presence or not of GlcN (added at T0h) and blood smears were realized regularly at ring, trophozoites and schizont stages.

PfEIF3i-HARibo parasites were cultured in the presence or absence of glucosamine in order to induce EIF3i protein knockdown. EIF3i protein levels were quantified by western blot. The two transgenic parasite clones show up to 80% reduction in EIF3i expression after glucosamine addition, indicating a successful knock-down of the target protein (Fig. 3C).

### PfEIF3i is essential for parasite growth and localizes to cytoplasm

To characterize the biological impact of EIF3i on parasite erythrocytic development, *P. falciparum* NF54 and PfEIF3i-HARibo lines were synchronized at the ring stage and the parasitemia was measured for two cycles by flow cytometry after or without glucosamine addition. EIF3i knock-down parasites (+ glucosamine) grow at a slower rate when compared to NF54 and to EIF3i-KD parasites cultured without glucosamine (Fig. 3D). The growth defect is also observed by Giemsa-stained thin blood smear in synchronized parasites. PfEIF3i-HARibo 40h-aged parasites cultivated with glucosamine show a delay in trophozoite maturation (Fig. 3E). Therefore, EIF3i seems to be essential for parasite intra-erythrocytic development. The protein is expressed at all intra-erythrocytic stages but mostly in trophozoites and schizonts (Fig. 4A). EIF3i localization in *P. falciparum* parasite was assessed by immunofluorescence (IFA), using anti-HA antibodies (Fig. 4B). The IFA show a cytoplasmic localization of the EIF3i protein in ring, trophozoites and schizonts, in agreement with its localization in human cells^11,13^.

**Figure 4.**
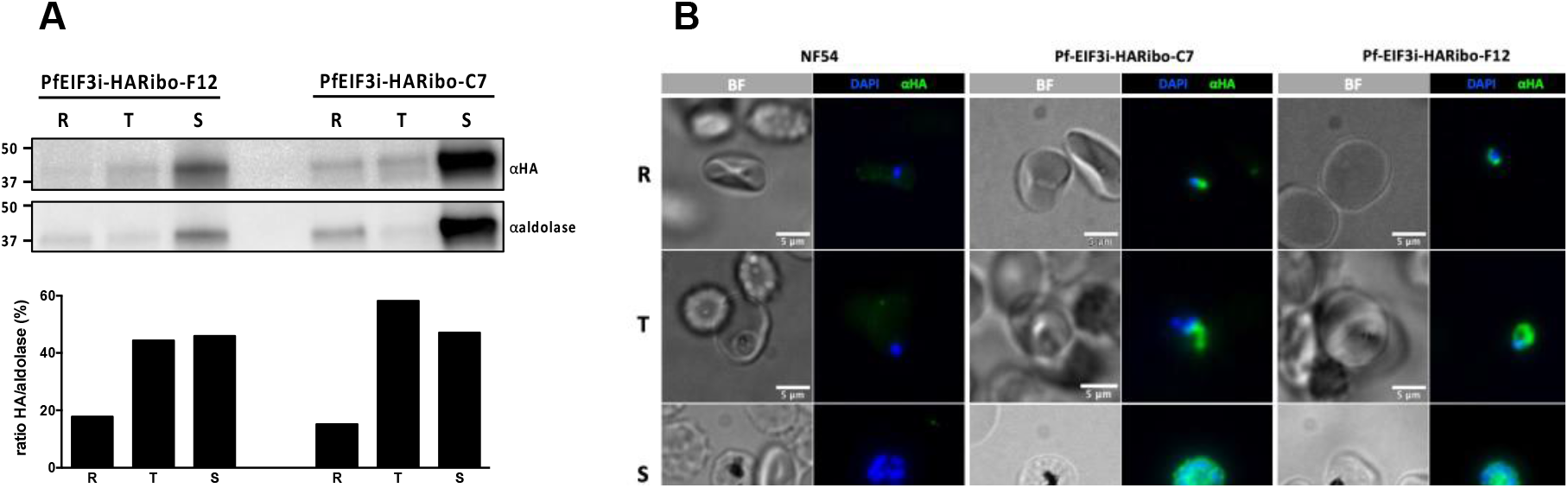
PfEIF3i is expressed at all intra-erythrocytic stages and localizes to the cytoplasm. **(A)** Extracts of PfEIF3i-HARibo clones C7 and F12 were collected after saponin lysis from ring (R), trophozoite (T) and schizont (S). EIF3i protein levels were quantified by western blot using anti-HA monoclonal antibodies and anti-aldolase antibodies as loading controls. Graph represents the ratio of HA/aldolase band intensities expressed in percentage. **(B)** Immunofluorescence assays of fixed RBCs infected with *Plasmodium falciparum* NF54 or PfEIF3i-HARibo parasites clones C7 and F12 at ring (R), trophozoite (T) and schizont (S) stages. EIF3i-HA was detected using anti-HA antibodies (green) and DNA was stained with DAPI (blue).

## Discussion

To decipher the mode of action of the quinoline-quinazoline-based inhibitors that were identified as potent new antimalarials^3^, we applied the TPP^8,16,17^ technique and identified potential targets of the bisubstrate compound **70** in asexual blood stages of *P. falciparum*. The eukaryotic translation initiation factor 3 subunit I (EIF3i) showed the highest score among the proteins stabilized by the inhibitor and thus chosen for further validation. We engineered a transgenic *P. falciparum* line expressing PfEIF3i tagged with an HA-tag to confirm that PfEIF3i is stabilized by compound **70** by CETSA-WB. Noteworthy, PfEIF3i has never been described as a primary drug target and has never been characterized in *Plasmodium*. To further study its function in the parasite, we generated an inducible knock-down of PfEIF3i. As predicted^18^, the protein is essential for the survival of the parasite as it blocks parasite development at the trophozoite stage when the protein is knocked-down by approximately 80%. PfEIF3i is expressed in the different stages of *P. falciparum* asexual blood cycle, which is coherent with the cell-cycle activity that we described for compound **70**^3^. Also, based on the mass-spectrometry datasets available in the *Plasmodium* database (PlasmoDB), plasmodial EIF3i is expressed in all life cycle stages of *P. falciparum*: asexual stages^19^, gametocytes^20^, and mosquito stages^21^ (ookinetes, oocyst, sporozoite). This protein is well conserved in eukaryotes and the EIF3 complex plays a key role in translation initiation and termination^22^.

In mammals, this large complex (approximately 800 kDa) is composed of 13 subunits, EIF3a-m^23^. Assembled together the subunits participate in several steps of translation initiation, termination ribosomal recycling, and in the stimulation of stop codon readthrough^24–27^. In PlasmoDB (plasmodb.org), only 12 EIF3 subunits are annotated but none have been studied yet. The PfEIF3 complex might have a role in drug-resistance^10^: in mefloquine-resistant *P. falciparum* strains, PfEIF3 was shown to be upregulated in a protein profiling analysis by mass spectrometry^28^. *Leishmania infantum* resistance to amphotericin-B was also associated with higher expression levels of EIF3^29^. PfEIF3i was also found in a mass spectrometry-coupled Cellular TSA (MS-CETSA) as a potential hit target of chloroquine, even though it was not identified in the labeled activity-based protein profiling (ABPP) used in parallel^30^. As bisubstrate inhibitors share a quinoline moiety in their scaffold with chloroquine, finding a common target protein is not surprising.

In conclusion, it is important to identify the targets of compounds that are potent antimalarials to use them in combination and anticipate and monitor resistance during further deployment^31^. Here we report a novel antimalarial target of the quinazoline-quinoline bisubstrate inhibitor **70** the protein PfEIF3i, which is essential for the parasite development and is expressed throughout the intraerythrocytic stages of *P. falciparum*, and during the different stages of the malaria life cycle, thus representing an attractive drug target.

## Methods

### Thermal Proteome Profiling

Experiments were carried out as previously described^16^ with minor modifications. Briefly, *P. falciparum* trophozoite lysate was exposed to varying concentrations of compound **70** (100µM-0.25nM) and a DMSO control for 3min, followed by thermal challenge at 37°C, 51°C or 57°C. Remaining soluble protein was analyzed by quantitative Mass Spectrometry. Data processing was carried out in R environment, using mineCETSA package 1.1.1. Proteins exhibiting drug dose dependent change in stability were identified using a set of criteria, including ≥PSM, R^2^ ≥0.8, ΔAUC ≥3*MAD.

### Plasmid constructs

The pSLI-HA-FKBP-T2A-yDHOH-Ribozyme plasmid was generated in our laboratory from the pSLI plasmid^32^ by In-fusion® (Takara) insertion of five PCR fragments coding for: 3x hemagglutinin epitope tag (3xHA), FKBP, T2A peptide, yeast dihydroorotate dehydrogenase (yDHODH), and *glmS* ribozyme sequences. The pSLI-EIF3i-HA plasmid was generated from the pSLI-HA-FKBP-T2A-yDHOH-Ribozyme plasmid by In-fusion® of two PCR fragments coding for the last 845 bp of *eif3i* cds (PlasmoDB^33^: PF3D7_0716800, primers P3F-P3R) and 3xHA (primers P4F-P4R) at NotI/XhoI restriction sites to replace FKBP, T2A, yDHODH and *glmS* ribozyme sequences. The pSLI-EIF3i-HARibo plasmid was generated from the pSLI-HA-FKBP-T2A-yDHOH-Ribozyme plasmid by In-fusion® of two PCR fragments coding for the last 845 bp of *eif3i* cds (primers P3F-P3R) and 3xHA (primers P5F-P5R) at NotI/SalI restriction sites to remove the FKBP sequence. All PCR were performed using KAPA HiFi DNA Polymerase (Roche 0795884600) with primers described in supplementary table 1. The last 845 bp of *eif3i* were PCR amplified from genomic *P. falciparum* NF54 DNA. The stop codon was removed from the original sequence and added after the 3xHA tag so it can be expressed with the *eif3i* gene. The final constructs are depicted in Figure 2A and 3A and primers used in this study are described in the supporting information (Table S1).

### Parasite culture and transfection

*P. falciparum* parasites were cultured using a standard protocol^34^. The laboratory strains used was NF54. Parasite transfections were done by electroporation of ring stages using 50 µg of purified plasmid NucleoBond Xtra kit (Macherey Nagel) as described elsewhere^35,36^. Drug selection was done using 2.66 nM of WR99210 (Jacobus Pharmaceuticals) for 5 days after transfection. After homologous recombination parasites were cloned by serial dilution^37^. PfEIF3i knockdown was achieved upon addition of 2.5 mM of glucosamine (Sigma, G1415).

### Western blot

Parasites were harvested using 0.15% saponin lysis of infected RBC (iRBC) followed by PBS washes. Proteins of the parasite pellets were extracted using lysis buffer (1 mM DTT, 2X Laemmli buffer (Bio-rad, 161-0737) and protease inhibitor cocktail (Roche, 11836170001) dissolved in PBS) and sonicated for 5 min. The equivalent of 1.10^7^ parasites were loaded per lane in a 4–20% Mini-PROTEAN® TGX Stain-Free™ gel (Bio-rad, 4568096) and proteins were separated and transferred onto a nitrocellulose membrane using the transblot semi-wet transfer system (Biorad). Blocking and antibody dilutions were performed in PBS-Tween-20 with 5% skim milk. Anti-HA (Abcam, ab9110), HRP-anti-*Plasmodium* aldolase (Abcam, ab38905) and anti-H3 (Abcam, ab1791) were used at 1:1000; secondary anti-rabbit HRP antibody was diluted at 1:5000. The blot was revealed using Super Signal West-Femto chemiluminescent substrate (Thermo Fisher Scientific).

### CETSA-Western Blot

Parasite total protein extracts were snap-frozen and stored at −80°C until further use. Approximately 3x10^8^ parasites diluted in 1ml of PBS were treated with 30 µM of compound **70**^3^ or pyrimethamine and the equivalent volume of DMSO for 1h at RT. Then the extracts were divided into 10 tubes of 100 µl and exposed to a 10-point temperature challenge from 40°C to 67°C with a 3°C fold increase, for 5 min. Following 3 min at RT, tubes were centrifuged at 15,000 g for 20 min at 4°C. Soluble proteins contained in the supernatant were separated and mixed with 4x Laemmli sample buffer (Bio-Rad, 161-0747) before being loaded on a SDS-PAGE gel (4–20% Criterion™ TGX Stain-Free™ Protein Gel, Bio-Rad #5678095). Western blot was then performed as described in the previous section.

### Growth curve

Parasites were synchronized at ring stage and incubated with fresh red blood cells at a starting parasitemia of 0.2%. Each day 20µl of the culture was collected and fixed with 0.025% glutaraldehyde and quenched after 1h with 15mM of ammonium chloride (NH_4_Cl). Fixed cells were then stained with 2X SYBR^®^ Green I (Lonza, 50512) and parasitemia was determined by flow cytometry using the Guava^®^ EasyCyte™ HT flow cytometer (Luminex).

### Immunofluorescence

iRBCs were fixed with 4% paraformaldehyde (EMS 15714) and 0.0075% glutaraldehyde (EMS 16220) in PBS for 30 min, as described previously (Tonkin et al., 2004). After a PBS wash, cells were permeabilized with 0.1% Triton-X 100 for 10 min and free aldehyde group were quenched with 50 mM NH_4_Cl for 10 min. Then cells were blocked with 1% bovine serum albumin (BSA) (Sigma A4503-50G) in PBS for 30 min and incubated overnight at 4°C with 1:1,500 anti-HA high affinity (Roche, 3F10). After three PBS washes, goat anti-rat Alexa Fluor 488 (Invitrogen, A-11006) secondary antibody was used at 1:2,000 with DAPI (FluoProbes FP-CJF800, 1 μg/mL). Finally, cells were mounted with VectaShield® (Vector Laboratories, H-1000) on glass slide. Images were acquired on a Deltavision Elite imaging system (Leica) and processed using ImageJ-FiJi software^38^.

## Author Contributions

The manuscript was written through contribution of all authors. All authors have given approval to the final version of the manuscript.

## Acknowledgements

This work was supported by Institut Pasteur-Institut Carnot (S-CR18089-02B15 DARRI CONSO INNOV 46-19; S-PI15006-10A INNOV 05-2019 ARIMONDO IARP 2019-PC), Pasteur Transversal Research Program (PTR 233-2019 HALBY), Pasteur Swiss Foundation grant, Agence Nationale de la Recherche (ANR EpiKillMal), and Pasteur-Roux-Cantarini Fellowship. Zbynek Bozdech’s contribution to the work was funded by the Singapore Ministry of Education grant number MOE-T2EP30120-0015, Jerzy Dziekan was funded by a NTU-PPF-2019 grant, and Peter Preiser contribution was supported by the National Research Foundation (Singapore) grant NRF-CRP24-2020-0005. Elie Hammam travel to Singapour was founded by the Merlion Grant.

The authors would like to thank Jessica Bryant of the Host Parasite Interactions unit at the Institut Pasteur for the pSLI-HA-FKBP-T2A-yDHOH-Ribozyme plasmid.

## Abbreviations

ABPP: activity-based protein profiling
ACTs: artemisinin-based combination therapies
AUC: area under the curve
cds: coding sequence
CETSA: cellular thermal shift assay
DNMT: DNA methyltransferase
EIF3: eukaryotic translation initiation factor 3
GlcN: glucosamine
HA: hemagglutinin
iRBC: infected RBC
ITDR: isothermal dose response
MAD: median absolute deviation
MS-CETSA: mass spectrometry-coupled cellular thermal shift assay
NH_4_Cl: ammonium chloride
PlasmoDB: *Plasmodium* database
TPP: thermal proteome profiling
yDHODH: yeast dihydroorotate dehydrogenase

## Supporting information

Table S1 references the PCR primers used in this study

---

For Table of Content Only

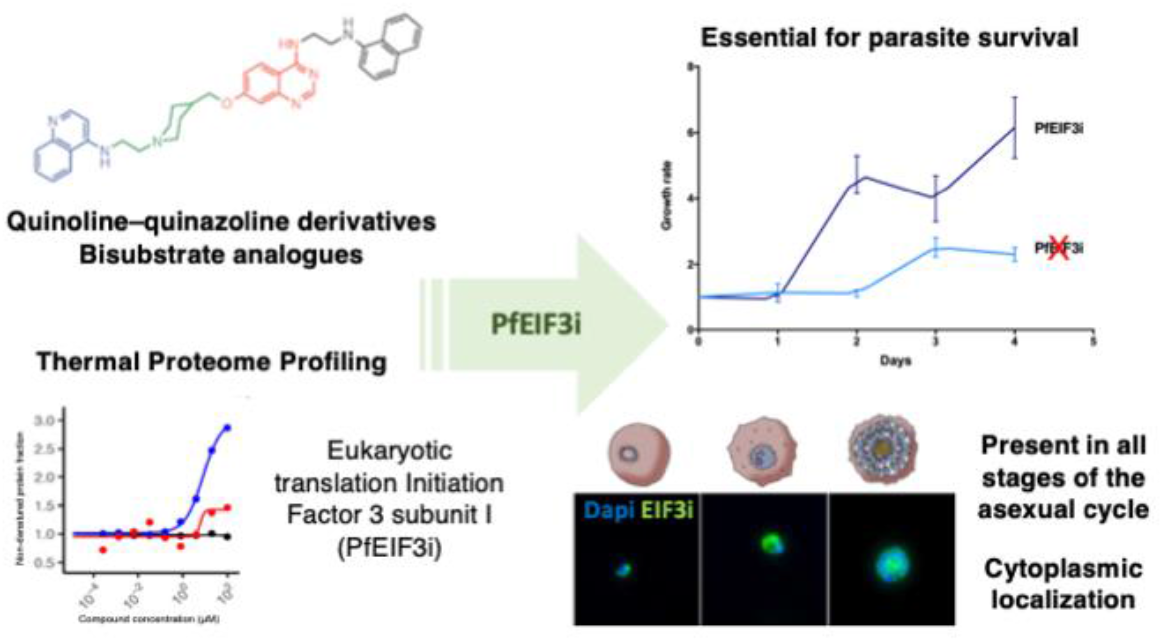

