## Supplemental Table 1 for "Quinazoline-quinoline bisubstrate inhibitors target eukaryotic translation initiation factor 3 in *Plasmodium falciparum*"

Table S1: Primers used in this study.

| Primer name | Sequence 5’ → 3’ |
|---|---|
| P1 | GATTTGCTATTTACAACAGGAAGGG |
| P2 | CACCCCGTGAGCATAATCCG |
| P3F | TAGAATACTCAAGCTGCGGCCGCTAAGATTATATGAATGTAGTGGGGCTGTG |
| P3R | AGGGTATCCACCGCCCCTAGGATCATATTTTCCTATAAAATAATCATTGTCAAAATGATAAATTC |
| P4F | TTTATAGGAAAATATGATCCTAGGGGCGGTGGATACCCTTAC |
| P4R | CATTAAGCTGCCATATCCCTCGAGTTACACCCCGTGAGCATAATCCGG |
| P5F | TGATCCTAGGGGCGGTGGATACCCTTAC |
| P5R | CTCCGTCGACCACCCCGTGAGCATAATCCGGAACG |
